# Nanopore and Illumina Sequencing Reveal Different Viral Populations from Human Gut Samples

**DOI:** 10.1101/2023.11.24.568560

**Authors:** Ryan Cook, Andrea Telatin, Shen-Yuan Hsieh, Fiona Newberry, Mohammad A. Tariq, Dave J. Baker, Simon R. Carding, Evelien M. Adriaenssens

## Abstract

The advent of viral metagenomics, or viromics, has improved our knowledge and understanding of global viral diversity. High-throughput sequencing technologies enable explorations of the ecological roles, contributions to host metabolism, and the influence of viruses in various environments including the human gut microbiome. However, the bacterial metagenomic studies frequently have the advantage. The adoption of advanced technologies like long-read sequencing has the potential to be transformative in refining viromics and metagenomics.

Here, we examined the effectiveness of long-read and hybrid sequencing by comparing Illumina short-read and Oxford Nanopore Technology (ONT) long-read sequencing technologies and different assembly strategies on recovering viral genomes from human faecal samples.

Our findings showed that if a single sequencing technology is to be chosen for virome analysis, Illumina was preferable due to its superior ability to recover fully resolved viral genomes and minimise erroneous genomes. While ONT assemblies were effective in recovering viral diversity, the challenges related to input requirements and the necessity for amplification made it less ideal as a standalone solution. However, using a combined, hybrid approach enabled a more authentic representation of viral diversity to be obtained within samples.

**Impact Statement:** Viral metagenomics, or viromics, has revolutionised our understanding of global viral diversity however long-read and hybrid approaches are not yet widespread in this field. Here, we compared the performance of Illumina short-read and Nanopore long-read assembly approaches for recovering fully resolved viral genomes from human faecal samples. We highlight Illumina’s short-read sequencing for recovering fully resolved viral genomes, while acknowledging Oxford Nanopore Technology’s long-read sequencing for capturing broader viral diversity. However, a hybrid approach, utilising both technologies, may mitigate the limitations of one technology alone.

**Data Summary:** All reads used in this study are available on European Nucleotide Archive (ENA) within the project PRJEB47625.

## Introduction

The study of uncultivated viruses through viral metagenomics, hereafter viromics, has shaped our understanding of global viral diversity. It is now more than 20 years since the sequencing of the first virome with the revolutionary linker-amplified shotgun library (LASL) approach [1]. Since then, the advent of high-throughput sequencing has driven a viromics revolution.

Viromics has uncovered ecological roles of viruses in diverse environments, shed light on the contribution of phages to the metabolism of their hosts [2–4], uncovered viruses as key players in the human gut microbiome [5, 6], allowed for the construction of uncultivated genomes larger than any of those obtained from culturing [7, 8] which are now being used to inform taxonomy [9–11]. However, bacterial metagenomics has taken advantage of the use of long-read sequencing technology, most notably Oxford Nanopore Technologies (ONT) and Pacific Bioscience (PacBio) to study complex ecosystems and microbiomes. Long read and hybrid assemblies have advanced bacterial genomics and metagenomics, improving completeness of metagenome assembled genomes (MAGs) and in some cases identifying modified DNA [12–17]. Although bacteriophages, the most abundant viruses in a virome, have far smaller genomes than their hosts, they are not always easily resolved with short read sequencing alone. For example, phages of the ubiquitous *Pelagibacterales* possess genomes with high micro-diversity and/or genomic islands [18, 19] which are known to cause fragmentation during assembly [20–23]. Long reads have the potential to cover the entire length of a viral genome, thus offering a solution to these issues.

There are a few notable examples of long-read and hybrid sequencing approaches being used for viromics. A pooled and MDA-amplified PromethION library was used alongside nonamplified Illumina libraries in a study of agricultural slurry, with the addition of ONT reads increasing the recovery of viral genomes [24]. Similarly, a long-read LASL approach sequenced on a MinION and paired with Illumina sequencing, dubbed VirION and later improved for VirION2, has also increased the number and completeness of viral genomes from marine samples [25, 26]. Additionally, a hybrid assembly approach has improved the assembly completeness and quality of individual phage genomes belonging to *Przondovirus* [27], utilising similar methods to those suggested for high quality bacterial genome assemblies [28].

However, whilst long-read sequencing may aid in the recovery of viral genomes, it is not without limitations and challenges.

Despite continuous improvements as new library preparation kits and flow cells are developed, ONT sequencing has a higher error rate compared to Illumina [29–31]. These higher error rates impact protein prediction through the introduction of erroneous stop codons, leading to truncated proteins and inaccurate functional annotations [30]. This has been shown to impact on viral identification when protein annotations are used for prediction of viral genomes (e.g., VIBRANT) [32, 33]. Polishing ONT virome assemblies with corresponding Illumina libraries can reduce this error rate and increase predicted protein lengths, although still not to the levels of Illumina-only assemblies [24, 33]. Furthermore, the yields of DNA obtained from virome extractions are typically low and the large input requirements of ONT sequencing can be prohibitive from samples of some environments (typically 1 µg of DNA in 50 µl of buffer). To overcome the input requirements, one of the first ONT virome studies utilised tangential flow filtration to concentrate large volumes of seawater [34], others have developed a LASL approach [25, 26], and MDA [24, 33]. However, MDA introduces biases into metagenomic libraries that can lead to the over-representation of viruses with small, circular ssDNA genomes [35–37]. Previously, the issues of MDA were overcome by pairing MDA-amplified ONT libraries with corresponding nonamplified Illumina libraries [24]. As the strengths and limitations of ONT sequencing are the opposite of those for Illumina, a hybrid approach may allow for the mitigation of both platform’s limitations.

Whilst there have been limited comparisons of long and short read sequencing for viromes, a recent study benchmarked sequencing technologies and assembly algorithms using a mock community of 15 phage genomes [33]. This work concluded that, as a single technology, Illumina performed best [33]. However, the addition of long reads (particularly ONT) increased recovery of viral genomes [33]. Here, we sought to investigate the impact of sequencing platform without amplification and assembly strategy on the recovery of viral genomes from human faecal samples and to offer recommendations for virome analyses.

## Materials and Methods

### Sample Collection, Processing and Sequencing

Sample collection, virome preparation, and Illumina sequencing were performed as part of a previously described comparison of PCR versus PCR-Free DNA library preparation for human faecal viromes [38]. The ethics for the previous study were approved by the University of East Anglia (UEA) Faculty of Medicine and Health Sciences (FMH) Research Ethics Committee (FMH20142015-28), Norwich in 2014, and by the Health Research Authority (HRA) NRES Committee (17/LO/1102; IRAS: 218545) in 2017. In brief, three human faecal samples were collected from healthy adult male donors. Samples were homogenised in sterile TBT buffer, centrifuged at 11,200 × g for 30 min at 10°C for two rounds, and filtered sequentially through 0.8 µm and 0.45 µm PES cartridge filters. Filtrates were PEG-precipitated to concentrate virus- like particles (VLPs), followed by treatment with DNAse and RNAse to remove free nucleic acids. DNA was extracted following a standard Phenol:Chloroform:Isoamyl alcohol protocol. To gain enough material for ONT sequencing, multiple DNA extracts were performed on aliquots of the same sample and then pooled. For full details, please see the previous publication [38].

Illumina libraries were sequenced using 2 × 150 bp paired-end chemistry (PE150) on the Illumina HiSeq X Ten platform [38]. This study uses the PCR-free Illumina libraries described in the previous publication [38]. ONT libraries were prepared using kit SQK-LSK109 and sequenced on a MinION with r9.4.1 flow cells. ONT basecalling was performed with Guppy v6.4.6.

### Quality Control of Reads

Illumina reads were trimmed with BBTools v39.01 following a previously reported protocol [39]. Reads were initially trimmed of adapters with bbduk.sh using ‘ktrim=r minlen=40 minlenfraction=0.6 mink=11 tbo tpe k=23 hdist=1 hdist2=1 ftm=5 ref=adapters.fà, followed by quality trimming with bbduk.sh using ‘maq=8 maxns=1 minlen=40 minlenfraction=0.6 k=27 hdist=1 trimq=12 qtrim=rl’, and error corrected with tadpole.sh using ‘mode=correct ecc=t prefilter=2’ (https://jgi.doe.gov/data-and-tools/software-tools/bbtools/). ONT reads were filtered using Filtlong v0.2.1 using ‘--min_length 500 --keep_percent 90’ (https://github.com/rrwick/Filtlong). Reads were inspected before and after trimming with FastQC v0.11.8 for Illumina reads (https://www.bioinformatics.babraham.ac.uk/projects/fastqc/), and NanoPlot v1.41.6 for basecalled ONT reads [40]. Read length distributions were summarised using the ‘stats’ command as part of SeqFu v1.17.1 [41]. For all assembly combinations, four assemblies were attempted; one for each of the three libraries and an additional co-assembly which pooled the individual libraries.

### Short-read and Binning Assemblies

Illumina reads were assembled using MEGAHIT v1.2.9 with ‘--k-min 21 --k-max 149 --k-step 24 --min-contig-len 1500’ [42]. Three virus-specific binning approaches were used on the MEGAHIT assembly. (1) vRhyme v1.1.0 was used with fastq files as input using bowtie2 v2.5.1 for read mapping [43], and the ‘--keep_bam’ flag was used to retain BAM files for use in another binning tool VAMB [44]. (2) VAMB v4.1.1 was used with default parameters [45]. (3) PHABLES v0.2.0 was used with default parameters [46]. For the “pooled” vRhyme and VAMB assemblies, the contigs were taken from the co-assembly but individual libraries were used for reads/BAMs. For downward processing, bins were concatenated into single contigs and fused with a single N using fuse.sh from BBTools v39.01 (https://jgi.doe.gov/data-and-tools/software-tools/bbtools/). All “binning” assemblies were combined with Illumina contigs native from their respective MEGAHIT assembly.

### Long-read and Hybrid Assemblies

ONT reads were assembled using a variety of assemblers, all of which were performed separately on “raw” reads and those processed with Filtlong. Canu v2.2 was used with ‘corMinCoverage=0 corOutCoverage=all corMhapSensitivity=high correctedErrorRate=0.105 genomeSize=5m corMaxEvidenceCoverageLocal=10 corMaxEvidenceCoverageGlobal=10 oeaMemory=32 redMemory=32 batMemory=200’ [47]. Flye v2.9.2-b1786 was used with ‘-- nano-hq –metà [15]. Redbean (wtdbg2) v2.5 was used with ‘-p 21 -k 0 -AS 4 -K 0.05 -s 0.05 - L 1000 --edge-min 2 --rescue-low-cov-edges’ [48]. Raven v1.8.1 was used with default parameters [49]. Unicycler v0.5.0 (using miniasm and Racon) was used with default parameters with ONT reads only, as well as hybrid assemblies that used both Illumina and ONT reads from their respective libraries [50–52].

All ONT assemblies underwent four rounds of polishing with Medaka v1.7.3 which used minimap2 v2.24-r1122 for alignments [53] (https://github.com/nanoporetech/medaka). To produce additional short-read polished assemblies, all medaka polished assemblies underwent one round of polishing with Polypolish v0.5.0 as described in the Polypolish documentation (https://github.com/rrwick/Polypolish). Firstly, reads from respective Illumina libraries were mapped using bwa mem v0.7.17-r1188 [54]. Alignments were filtered using polypolish_insert_filter.py, and Polypolish was then used with default parameters [55].

### Viral Identification and ORF Prediction

Viruses were predicted from the assemblies using geNomad v1.5.2 [56]. Predicted viruses were subsequently processed using CheckV v1.0.1 that uses db v1.5 [57]. Open reading frames (ORFs) were predicted using Prodigal-gv v2.11.0-gv, a fork of Prodigal [58] that has been optimised for viral gene prediction (https://github.com/apcamargo/prodigal-gv).

### Recovery of Viral Diversity

Predicted viruses with a CheckV completeness estimate ≥ 50% (medium-quality and above) were dereplicated to form viral operational taxonomic units (vOTUs) following Minimum Information about an Uncultivated Virus Genome (MIUViG) standards (95% ANI over 85% length) using BLAST v2.14.0+ alongside the anicalc.py and aniclust.py scripts described in the CheckV documentation (https://bitbucket.org/berkeleylab/checkv/src/master/) [57, 59, 60]. Translated proteins predicted using Prodigal-gv v2.11.0-gv were used as input for vConTACT2 v0.11.3 alongside the INPHARED database (July 2023) with ‘--min-size 1’ [61, 62]. Viral clusters (VCs) were treated as genera, with those containing no reference sequences being described as novel. Pooled Illumina reads were mapped to sequences for which the VC contained ONT assembled sequences but no Illumina-based assemblies using bbmap.sh BBTools v39.01 (https://jgi.doe.gov/data-and-tools/software-tools/bbtools/), to determine if the VC could be detected in the Illumina data without being assembled. Presence was defined as ≥ 1× coverage over ≥ 75% of contig length [23].

### CrAssphage Analysis

Sequences that clustered with CrAssphage LMMB (MT006214) were processed using Clinker v0.0.28 to compare genome synteny and completeness [63].

### Data Visualisation

Unless otherwise stated, all plots were produced in R v4.2.2 [64] using ggplot2 v3.4.2 [65]. The vConTACT2 network was visualised using Cytoscape v3.9.1 [66]. Figure 3A was produced using UpSetR v1.4.0 [67], and figure 3B was produced using Clinker v0.0.28 [63].

## Results

To compare the performance of commonly used assembly algorithms for recovery of viral genomes from faecal samples, we sequenced three human gut viromes using Illumina and ONT sequencing and tested a number of assembly strategies for reconstructing virus genomes from the read datasets (Supplementary Table 1).

### Data Generation

Illumina sequencing of nonamplified viromes was carried out on the HiSeq X Ten platform using 150 bp paired-end libraries. The Illumina libraries generated 5.59, 4.96 and 6.27 Gbp of data with a mean phred score ≥30 ranging from 92.37% to 94.35% (Supplementary Table 2). Post trimming and quality control, Illumina libraries were 5.48, 4.9 and 6.16 Gbp with a mean phred score ≥30 ranging from 92.64% to 95.73% (Supplementary Table 2). To generate enough material for ONT sequencing, multiple DNA extracts from aliquots of the same sample were pooled prior to sequencing on a MinION with r9.4.1 flow cells. ONT libraries generated 8.68, 5.1 and 5.43 Gbp data with median Q scores of 14.1, 13.1 and 13.3 respectively (Supplementary Table 2). Post filtering with Filtlong, ONT libraries were 7.73, 4.1 and 3.89 Gbp with median Q scores of 14.8, 13.4 and 13.9 respectively (Supplementary Table 2). Summaries of read length distributions for all libraries are shown in Supplementary Table 2.

### Virome Assembly

Assemblies were produced using a variety of strategies and assemblers, including hybrid and binning approaches (Supplementary Table 1). For each assembler combination, four assemblies were attempted, one for each library and an additional pooled library. Short-read assemblies were performed with MEGAHIT only, as assembly comparisons for Illumina reads have previously shown it to provide high-quality assemblies [23, 68]. All ONT assemblies were attempted using both “raw” reads as well as those filtered with Filtlong. Additionally, all ONT assemblies were polished with Illumina reads from their respective library using PolyPolish.

Out of a possible 104 assemblies, 93 were produced successfully (Supplementary Table 1). Sample 03 failed to assemble using VAMB due to not satisfying the minimum number of contigs required for input (≥ 4096). Unicycler failed to yield assemblies for sample 01 and the pooled sample when using ONT reads only, and the pooled sample when using ONT and Illumina reads together. The Unicycler assemblies failed both before and after filtering ONT reads with Filtlong. All assemblies were given a maximum of 88 cores and 1,500 Gb of memory on an Intel® Xeon® Gold 6238 CPU @ 2.10GHz node with four CPUs that have 22 cores each, the failure is therefore unlikely to be due to limited computational resources. However, as Unicycler was designed for single genomes rather than mixed community samples, this is not surprising.

Regarding total assembly size, the largest assemblies were obtained from Illumina data (20 – 125 Mb per assembly; Table 1; Supplementary Table 3). However, these assemblies were highly fragmented, containing the highest number of contigs (1,851 – 27,614 per assembly), with the shortest contig lengths (1.85 – 2.51 Kb median contig lengths per assembly). The total assembly size and median contig length for ONT-based assemblies varied greatly with the assembler used. Unicycler and Raven obtained the highest median contig lengths (22.42 –36.45 Kb median contig lengths per assembly), although this was at the cost of smaller total assemblies (7.97 – 49.4 Mb per assembly). Flye and wtdbg2 produced larger total assemblies (14.86 – 108.51 Mb per assembly), although with shorter median contig lengths (5.31 – 11.55 Kb median contig lengths per assembly). Unicycler with ONT and Illumina reads together, and Canu produced small assemblies (7.41 – 27.72 Mb per assembly) with relatively short contig lengths (6.57 – 20.1 Kb median contig lengths per assembly). The use of Filtlong prior to assembly had substantial impacts on some ONT assemblers, increasing median contig lengths from 26 to 32.5 Kb and 25.5 to 27 Kb for Unicycler (with ONT reads only) and Raven respectively. The same effects of Filtlong were not observed for Canu, Flye, wtdbg2, and Unicycler (with ONT and Illumina reads together). Unsurprisingly, polishing ONT assemblies with Illumina reads led to modest reductions in total assembly size.

**Table 1:** Summary Statistics of Virome Contigs and Contig Bins by Assembly Type.

| Type | Min Length (Kb) | Max Length (Kb) | Median (Kb) | Mean (Kb) | SD (Kb) | Min # Contigs | Max # Contigs | Median # Contigs | Mean # Contigs | SD | Min Assembly (Mb) | Max Assembly (Mb) | Median Assembly (Mb) | Mean Assembly (Mb) | SD (Mb) |
| --- | --- | --- | --- | --- | --- | --- | --- | --- | --- | --- | --- | --- | --- | --- | --- |
| Bins | 1.5 | 2,568.45 | 2.25 | 6.51 | 33.44 | 1,851 | 27,595 | 9,443 | 10,539.27 | 7,001.78 | 20.15 | 124.97 | 69.75 | 68.62 | 40.02 |
| Hybrid | 1.5 | 688.79 | 7.44 | 20.57 | 42.03 | 352 | 1460 | 597 | 802.67 | 519.64 | 9.57 | 27.72 | 12.26 | 16.51 | 8.75 |
| Illumina | 1.5 | 385.38 | 2.42 | 4.62 | 9.46 | 3,559 | 27,617 | 12,319.5 | 13,953.75 | 10,188.24 | 20.15 | 124.61 | 56.65 | 64.51 | 44.9 |
| ONT | 1.5 | 1,301.1 | 10.45 | 23.62 | 47.41 | 166 | 6,154 | 694.5 | 1,399.42 | 1,502.44 | 7.41 | 108.51 | 20.88 | 33.05 | 27.16 |
| Polished | 1.5 | 1,300.91 | 10.45 | 23.61 | 47.4 | 166 | 6,154 | 694.5 | 1,399.42 | 1,502.44 | 7.41 | 108.48 | 20.88 | 33.04 | 27.15 |

### Recovery of Viral Genomes

To compare the performance of the assemblies in recovering viral genomes, we predicted viruses with geNomad and assessed their completeness with CheckV. For the purposes of this section of results, we included genomes with an estimated completeness of ≥ 50% (Medium-quality and above).

The choice of assembly algorithm used to assemble ONT reads had a substantial effect, as ONT assemblies performed both highest and lowest for viral genome recovery depending on assembler used (Figure 1A; Supplementary Table 4). The Flye assemblies obtained 108 to 459 genomes which was marginally increased after polishing with Illumina reads (111 – 464), representing the highest values of any assembly tested. Second to this were the ONT wtdgb2 assemblies that obtained 84 to 308 genomes prior to polishing with Illumina reads, and 83 to 313 after. Following this were the Illumina based assemblies with MEGAHIT obtaining 80 to 212 viral genomes, that was further increased using binning approaches VAMB (80 – 216), Phables (83 – 218), and vRhyme (88 – 228). This was followed by Raven (61 – 213), Unicycler with ONT reads (45 – 140), Canu (61 – 135), and Unicycler with ONT and Illumina reads together (71 – 123). Typically, there was little difference between ONT assemblies that had been pre-processed with Filtlong or not. However, there was a substantial difference for the ONT-only Unicycler assemblies (Figure 1A; Supplementary Table 4).

**Figure 1.**
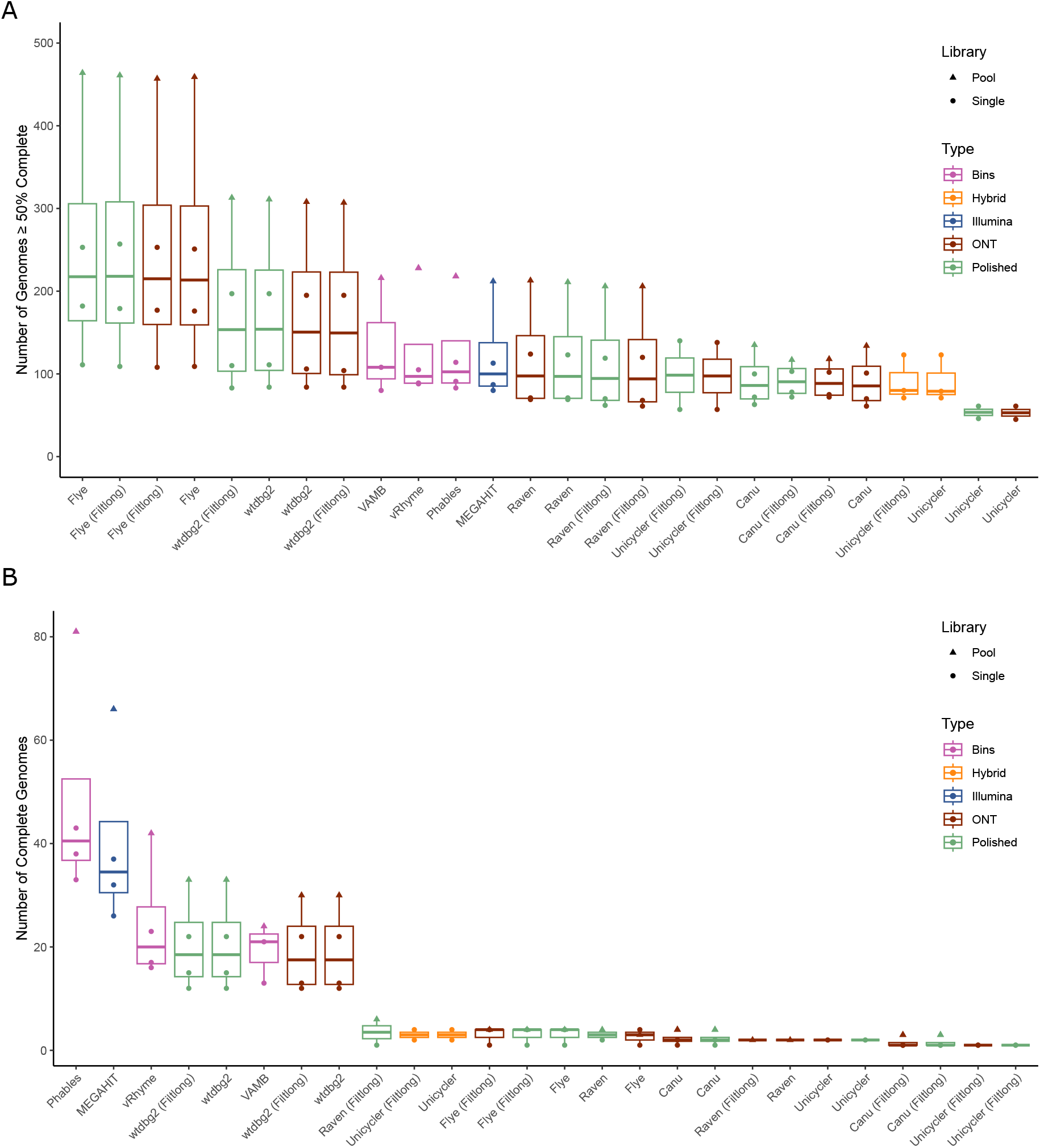
Recovery of Viral Genomes. The number of viral genomes per assembly predicted to be (**A**) ≥50% complete and (**B**) 100% complete.

Whilst ONT based assemblies recovered the highest number of genomes with ≥ 50% estimated completeness, Illumina assemblies obtained the most fully resolved viral genomes (100% complete; Figure 1B; Supplementary Table 4). The Illumina MEGAHIT assemblies resolved 26 to 66 complete genomes, and this was increased to 33 - 81 using Phables (Figure 1B; Supplementary Table 4). The ONT-only assemblies with the most complete genomes were produced using wtdbg2, obtaining 12 to 30 complete genomes, increasing to 12 to 33 after polishing with Illumina reads. All other long read assemblers performed poorly at recovering 100% complete viral genomes (Figure 1B; Supplementary Table 4).

### Assembly Quality Assessment

To assess the presence of potential errors in viral genomes, such as duplications, we used CheckV warning flags as indicators. These flags were raised either due to a high k-mer frequency or when a contig’s length exceeded 1.5 times the expected genome length but was not predicted to be an integrated prophage (Figure 2; Supplementary Table 4).

**Figure 2.**
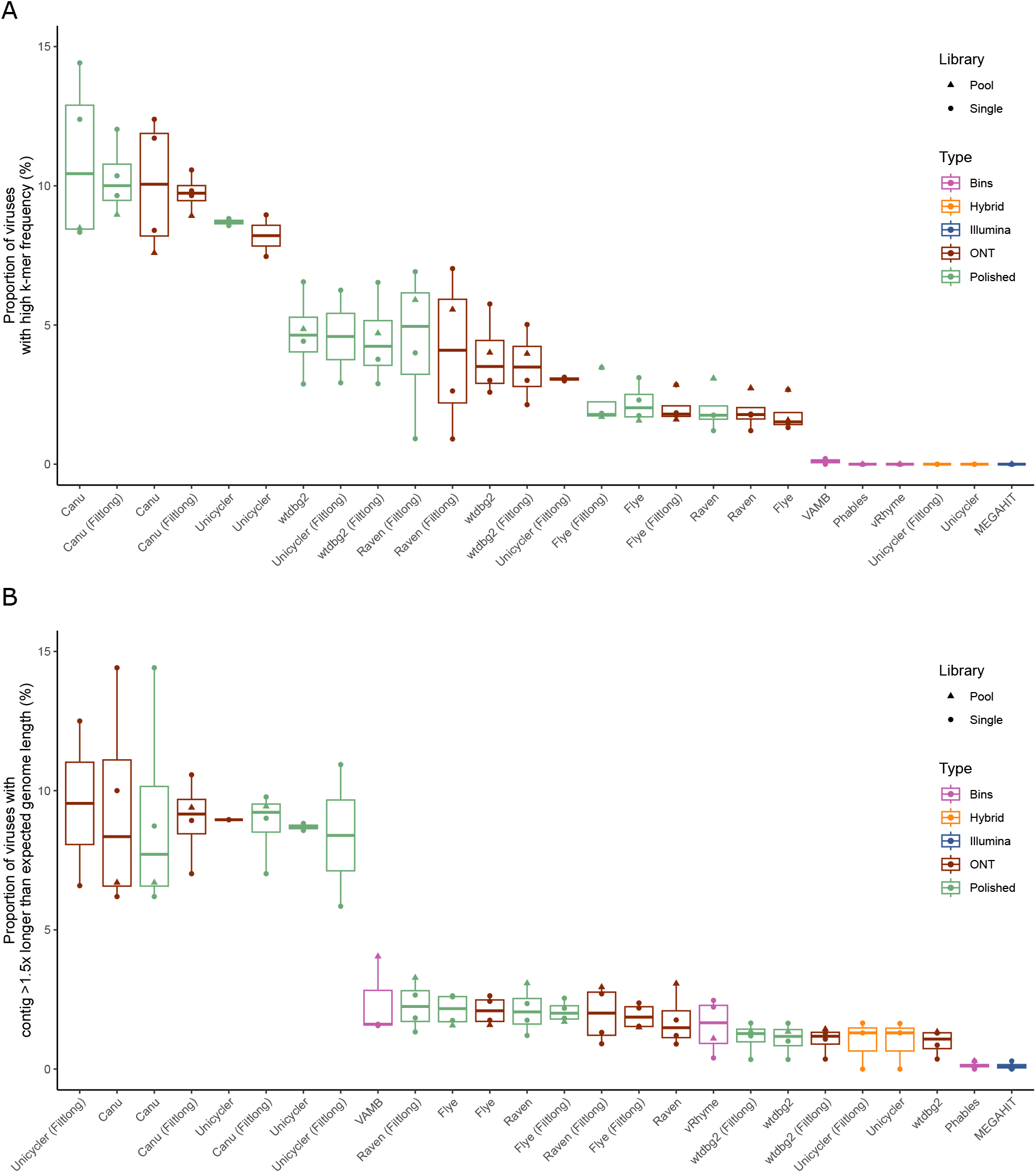
Frequency of Erroneous Viral Genomes. The number of viral genomes per assembly with CheckV warning flags for (**A**) high k-mer frequency and (**B**) contig exceeding 1.5 x expected genomes length without being predicted as a prophage.

Illumina assemblies produced using MEGAHIT had no sequences with a high k-mer frequency warning, those binned with Phables and vRhyme also had no sequences with this warning (Figure 2A; Supplementary Table 4). Binning with VAMB increased the frequency to 0.0 – 0.2% of predicted viruses with the warning. Similarly, when employing direct hybrid assemblies with both Illumina and ONT reads through Unicycler, no sequences were identified with high k-mer frequency issues (Figure 2A; Supplementary Table 4). Conversely, when focusing on ONT-based assemblies, the frequency of sequences with this warning ranged from 0.9 to 14.4 % per assembly, with large variation among the assemblers. The assemblies produced using Canu yielded the highest frequency (7.6 – 14.4%), followed by Unicycler (2.9 – 9.0%), wtdbg2 (2.1 – 6.6 %) Raven (0.9 – 7.0%), and Flye (1.3 – 3.5%; Figure 2A; Supplementary Table 4). Notably, polishing the ONT assemblies with Illumina reads typically increased the occurrence of this warning message; this may be due to the removal of errors in the ONT assembly, causing the two duplicated regions to become more similar as the errors are removed (Figure 2A; Supplementary Table 4).

Similarly, Illumina assemblies produced using MEGAHIT obtained the lowest number of sequences flagged as being >1.5 times longer than the expected length (0.0 – 0.3%; Figure 2B). Those binned with Phables also obtained 0.0 – 0.3% (Figure 2B; Supplementary Table 4). However, when using other binning approaches, the occurrence of this warning increased. The vRhyme assemblies ranged between 0.4 – 2.5%, and VAMB between 1.6 – 4.0% (Figure 2B; Supplementary Table 4). Assemblies using only ONT reads had a range of 0.4 – 14.4% per assembly for this warning, with variation depending on the assembler used (Figure 2B; Supplementary Table 4). Among the ONT-only assemblies, Canu had the highest frequency (6.2 – 14.4%), followed by Unicycler (5.8 – 12.5%), Raven (0.9 – 3.3%), Flye (1.5 – 2.6%), and wtdbg2 with the lowest (0.3 – 1.7%; Figure 2B; Supplementary Table 4). The levels obtained by direct hybrid assemblies produced using Unicycler with Illumina and ONT reads together were similar to the best performing ONT-only assemblies (0 – 1.7%; Figure 2B; Supplementary Table 4). No clear pattern was observed regarding the use of Filtlong before assembly and/or the polishing process with Illumina reads after assembly in relation to this particular metric, for some assemblers the frequency increased, for others it decreased (Figure 2B; Supplementary Table 4).

### Differences in Predicted Viral Diversity

To determine any differences in viral diversity recovered within each assembly, phage contigs with a predicted completeness of medium and above were clustered to form vOTUs (95% ANI over 85% alignment; approximate species), which were then clustered using vConTACT2 alongside INPHARED (July 2023) to form viral clusters (VCs) that are approximately analogous to genera/subfamily level taxonomy. We examined the number of VCs per assembly type to quantify the estimated viral diversity.

The ONT assemblies recovered 453 VCs across the three donor samples, with 185 of these being exclusive to ONT based assemblies (Figure 3A; Supplementary Table 5). However, the number of VCs varied with assembler used. Flye assemblies obtained the highest number of VCs with 97 to 349, followed by wtdbg2 (76 – 278), Raven (59 – 202), Canu (42 – 87), and Unicycler (39 – 59; Figure 3A; Supplementary Table 5). These predictions changed very little after polishing with Illumina reads. The Illumina assemblies produced using MEGAHIT obtained 75 to 202 VCs, similar to Phables at 75 to 203 (Figure 3A; Supplementary Table 5). Interestingly, a slight reduction in the number of VCs recovered was observed following use of VAMB (74 – 171) and vRhyme (69 – 196; Figure 3A). It may be that fragmented Illumina contigs derived from the same genome were clustered into separate VCs when using the Illumina MEGAHIT assemblies, and that some of these were resolved into the same genome through binning. The direct hybrid assemblies produced using Unicycler with ONT and Illumina reads together obtained the fewest VCs (74 – 111), and only two of these were exclusive to these assemblies (Figure 3A; Supplementary Table 5).

**Figure 3.**
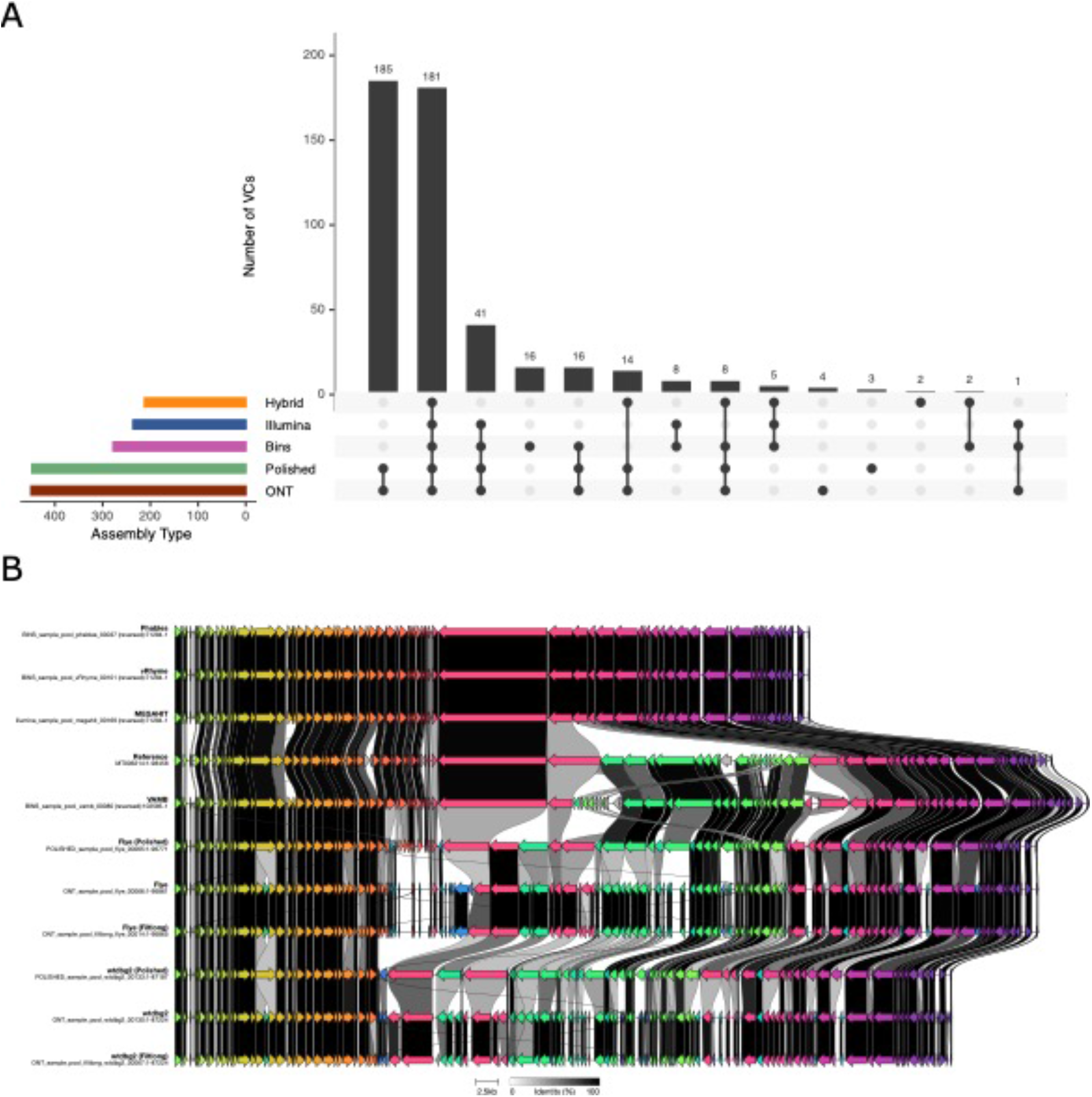
Recovery of Viral Clusters per assembly type. (**A**) The number of viral clusters recovered by each sequencing approach. (**B**) Comparison of synteny and genome architecture of virome sequences compared to CrAssphage LMMB (<u>MT006214</u>).

To determine if the ONT-only VCs were present in the Illumina libraries but not being assembled, we mapped reads to the sequences in these VCs to determine their presence. Of the 206 VCs that were exclusive to ONT/Polished/Hybrid assemblies, 107 contained sequences that were present in the Illumina libraries. The Illumina reads which mapped to the ONT-only clusters were then mapped back to the vOTUs obtained from the Illumina MEGAHIT assembly. Read mapping recruited 80.3% of reads to the MEGAHIT vOTUs, with 99 vOTUs obtaining ≥ 1× coverage. A large proportion of the ONT-only VCs therefore contained sequences similar to those in the Illumina-only assemblies. However, their forming of separate VCs is likely due to incorrect proteins predictions resulting from higher error rates and erroneous stop codons.

### Recovery of CrAssphage LMMB

We noticed that several assemblies contained near-complete genomes that were highly similar to CrAssphage LMMB (<u>MT006214</u>). As the study of *Crassvirales* is a common component of human virome analyses, we decided to examine these sequences in comparison to the CrAssphage LMMB to determine their completeness and synteny (Figure 3B).

The Illumina MEGAHIT sequence had high levels of nucleotide identity to that of CrAssphage LMMB with highly conserved genome architecture and synteny (Figure 3B). However, there was a ∼27 Kb segment of the genome that was missing from the MEGAHIT sequence. The use of vRhyme and Phables did not resolve this section, and their assembly was the same as that of MEGAHIT-only. Whereas binning with VAMB was able to recover the section that was missing from the MEGAHIT assembly, leading to the most complete genome of any assembly used (Figure 3B). Regarding ONT assemblies, Flye was able to resolve the full length of the genome, including the ∼27 Kb segment that was missing from the MEGAHIT assembly. Whilst wtdbg2 was able to resolve most of the genome, including the segment missing from MEGAHIT, a different ∼11 Kb section was missing from the wtdbg2 assemblies (Figure 3B). While they were able to recover the full CrAssphage genome, the ONT-only assemblies contained a high number of highly fragmented ORFs (Figure 3B). It is likely that the higher error rate associated with ONT sequencing impacted the ORF prediction on these sequences. Polishing the Flye and wtdbg2 sequences with Illumina reads led to longer ORFs that appeared to be more congruent with CrAssphage LMMB, although there was still a clear difference when compared to the reference genome and Illumina assemblies (Figure 3B).

### Impacts of Assembler on Predicted Protein Lengths

As higher error rates are associated with the introduction of erroneous stop codons within predicted proteins, resulting in their truncation, we examined the length of predicted proteins per assembly as a proxy for error rates.

The Illumina assemblies produced using MEGAHIT had a median translated ORF length of 142 amino acids (AAs), versus the ONT based assemblies with a median length of 127 AAs (Figure 4). Polishing the ONT assemblies with Illumina reads increased the median ORF length from 127 to 133 AAs (Figure 4). Although the polishing likely removed some erroneous stop codons, the ORF length was still lower than that of Illumina only assemblies (142 AAs; Figure 4). However, the increase of median translated ORF length post-polishing did vary with assembler used. Assemblies produced using wtdbg2 increased from 120 to 128 median AAs, Flye from 122 to 128 with Filtlong and from 123 to 128 without, Unicycler from 129 to 134 with Filtlong and from 136 to 143 without, Canu from 136 to 146 with Filtlong and from 134 to 145 without, and Raven from 137 to 142 with Filtlong and from 137 to 141 without (Supplementary Figure 1). Therefore, polishing ONT assemblies with Illumina reads did restore ORF lengths to those comparable with Illumina only assemblies for some assemblers tested, specifically Unicycler, Canu and Raven.

**Figure 4.**
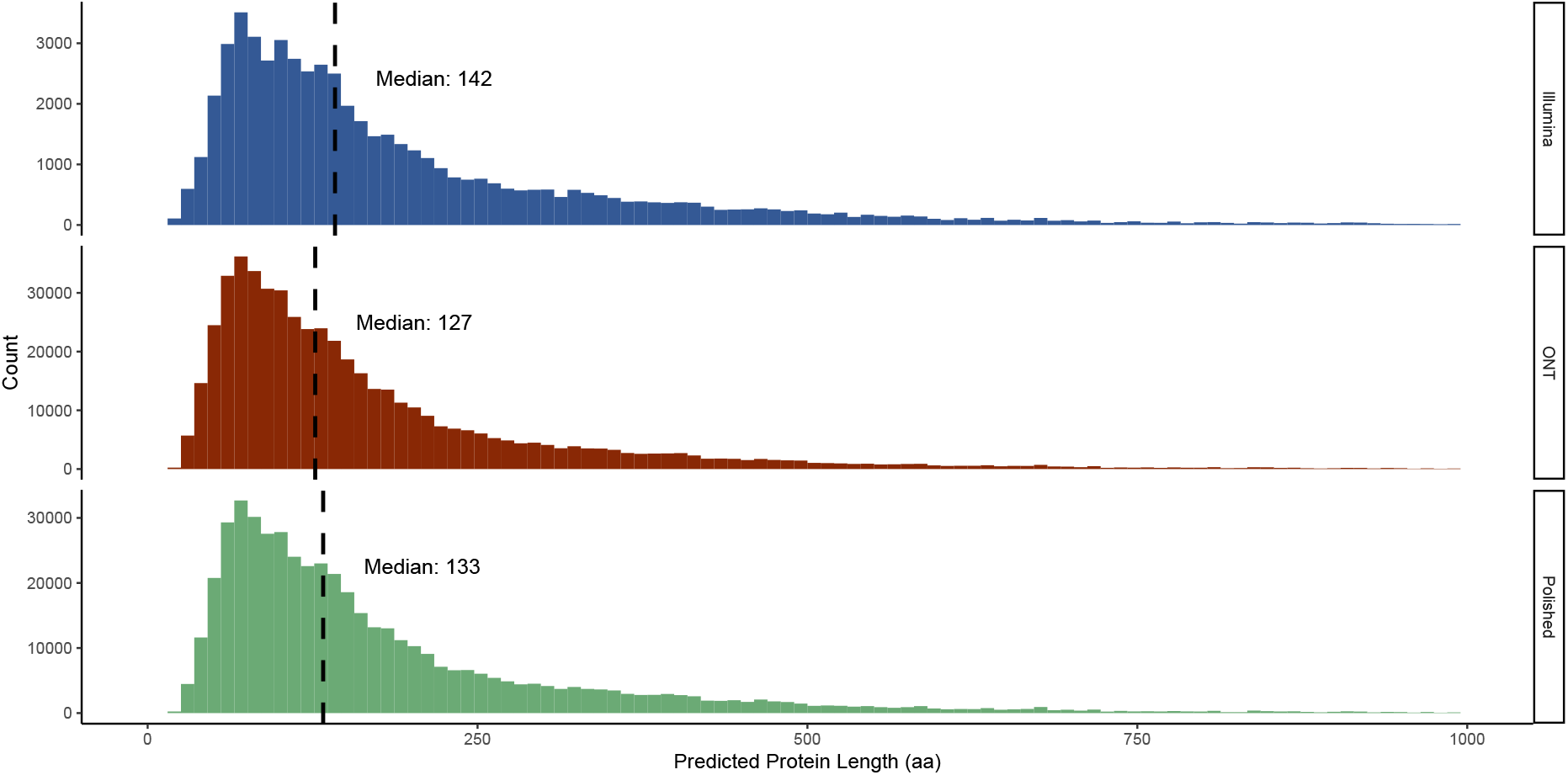
Effect of Polishing on Predicted Protein Lengths. Distribution of protein lengths (aa) with median value indicated by a dashed vertical line.

## Discussion

The use of long-read and hybrid sequencing approaches in metagenomics is becoming more common and has been demonstrated to increase the quality and completeness of bacterial genomes [12–17]. These approaches are beginning to emerge in the field of viromics, although there are still relatively few analyses of viromes sequenced with long-read and hybrid approaches. Currently, there are few comparisons of Nanopore and Illumina sequencing for the recovery of viral genomes using mock communities [25, 33]. However, there is little study into the recovery of viral genomes from complex natural samples using long-read sequencing. Here, we compared the performance of Illumina and Nanopore sequencing for the recovery of viral genomes from human gut samples using a number of assembly approaches.

Short-read assemblers have previously been benchmarked for recovery of viral genomes from mock communities, with metaSPAdes and MEGAHIT performing best [23, 68]. Whilst metaSPAdes is preferred for assembly quality, it is more computationally demanding than MEGAHIT [23, 68]. Hence, we used only one short read assembler: MEGAHIT [23, 68]. For long-read sequencing, we chose several commonly used assemblers; Flye, Canu, Raven, wtdbg2, and Unicycler. Unicycler was also used for a direct hybrid assembly with combined ONT and Illumina reads. Whereas Flye, Canu, Raven, and wtdbg2 have all been developed with options for metagenome assembly, Unicycler is optimised for assembling single genomes and is therefore expected to perform less well in viromics than the other assemblers.

Determining which sequencing technology and assembler combination worked best in this work was not trivial, and depends on the specific research question (Table 2). With the aim of recovering the most fully resolved viral genomes, our findings suggest Illumina is the best approach, whereas for maximising recovered viral diversity ONT is ideal (Table 2). However, previous study of Illumina sequencing for the recovery of viral genomes has uncovered false DTRs at the termini of Illumina sequences arising from repeated assembly artefacts, calling into question the validity of “complete” viral genomes recovered from Illumina-only assemblies [27]. Similarly, the increase in predicted viral diversity from ONT sequencing is not necessarily a clear benefit, as further analysis of the data brings the reliability of the predictions into question. We found that much of the increased diversity in the ONT data was likely the result of erroneous protein predictions, leading to incorrect clustering of contigs.

**Table 2:**
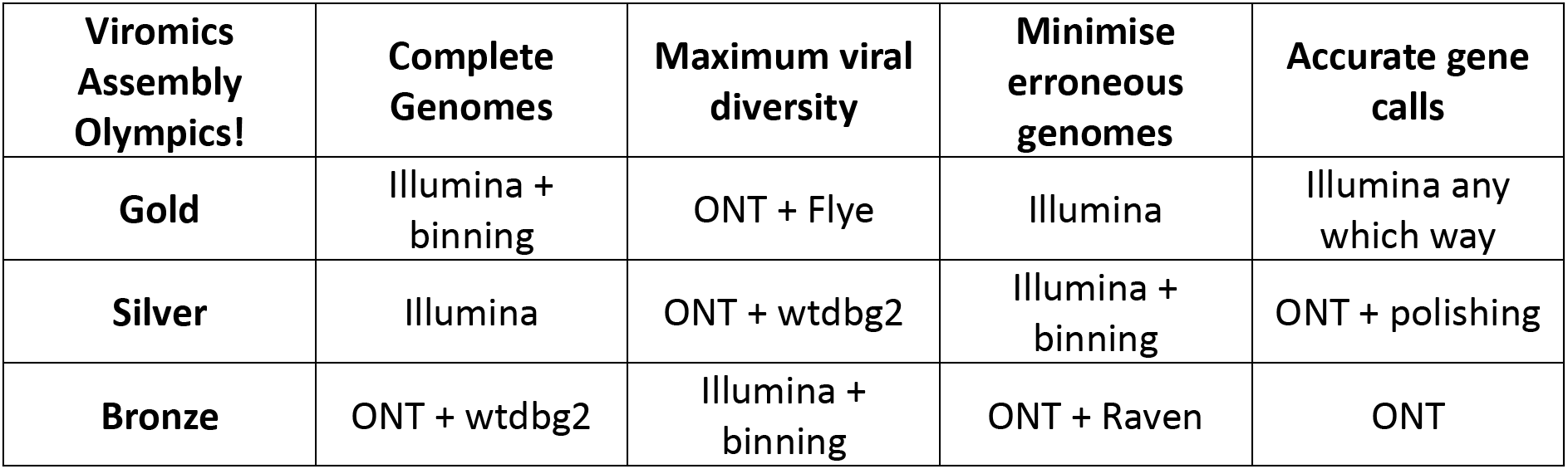
Relative Performance of Assembly Approaches for Common Objectives.

This finding was consistent with a previous comparison in which long-read assemblies were shown to vastly overestimate viral diversity within a mock community analysis [33]. Furthermore, there was also the associated cost of higher frequencies of erroneous genomes and higher error rates. However, there are reports of virus genomes not being recovered with Illumina sequencing whereas ONT could potentially provide a solution, even if more error prone [69, 70].

Therefore, taken altogether, we suggest that if only one sequencing technology were used for a virome analysis, Illumina should be preferred to ONT. Although the ONT assemblies performed favourably in recovering viral diversity, the input requirements and need for amplification are still the largest barrier for virome sequencing. However, long reads are not without their benefits. The use of hybrid approaches that combine Illumina sequencing with ONT may best capture viral diversity within a sample and overcome issues associated with amplification, although this increases costs in using more than one sequencing technology [24, 25, 33].

The continual development and improvement of ONT flow cells and assembly algorithms will improve the quality of assemblies. To date, there is relatively little optimisation of ONT sequencing specifically for viromics. For example, it is well documented that bacteriophages often have heavily modified DNA, and this may impact on the accurate basecalling of their genomes [71–74]. It was also previously reported that too much sequence depth is detrimental for ONT virome assembly [33], although downsampling prior to assembly will remove genomes of lower abundance. Furthermore, there are clear differences in ONT assemblers, with no single assembler performing best in all metrics. In bacterial genomics, the use of multiple assemblers can generate more trustworthy consensus sequences in the form of Trycycler [75]. There is currently no such approach for metagenomes and viromes. To accurately uncover more viral diversity from natural samples, there is a clear need for assembly algorithms and library preparation methods that are optimised for viral metagenomics.

On the laboratory side, the development of the VirION2 protocol which included optimisations in the choice of polymerase, PCR cycles and DNA shearing has led to increased read lengths and reduced the input DNA requirement to 1 ng [25]. Unfortunately, the laboratory work for this study was initiated before the VirION2 protocol was published and we therefore couldn’t incorporate some of the optimisations. Additionally, other emerging long-read technologies could be promising for viral metagenomics. Notably, PacBio HiFi sequencing is able to improve the completeness of bacterial MAGs and obtain far lower error rates than ONT sequencing [76].

Even with improvements that allowed for increased genome recovery from viral metagenomes, there are still large biases in the viruses recovered. This study, and the overwhelming majority of virome studies, focus exclusively on the DNA fraction and this is further biased towards dsDNA viruses. Recent mining of global metatranscriptomes has expanded currently known RNA viral diversity, and suggests that RNA viruses may be a critical but currently neglected component of the global virome [77]. There are virome protocols which include cDNA synthesis for the sequencing of RNA alongside DNA [78], however the resulting fragments may be too short to take advantage of long reads. Whilst direct RNA sequencing with ONT is available, the input requirements are prohibitive for most virome studies. There are still many technical challenges to capturing true viral diversity within an environmental sample.

## Conclusions

The use of viral metagenomics has accelerated our understanding of viral diversity within a plethora of environments, although much viral diversity remains unseen and continual improvements to library preparation methods, sequencing technologies, and assembly algorithms are needed to understand true viral diversity. Illumina based approaches may still offer the current gold standard for viromics, and the use of viral-specific binning algorithms, such as Phables, may aid the recovery of viral genomes. The addition of long reads uncovers additional viral diversity, although stringent quality control should be used to minimise the occurrence of erroneous viral genomes. As the choice of sequencing technology and assembly approach will impact the observed viral diversity, these methodological choices should always be considered in the interpretation of results.

## Acknowledgments

We would like to thank Dr. Oliver “Oli” Charity for his contributions to this work. Ollie was a talented bioinformatician who took lead on this work during data analysis, providing ideas and discussion that helped to shape the subsequent analysis.

## Author contributions

R.C, S.R.C and E.M.A conceived the study. S.Y.H, F.N., M.A.T carried out experiments and collected data. D.B and M.A.T prepared DNA libraries and performed sequencing. R.C, A.T and E.M.A performed the bioinformatic analysis. R.C, A.T and E.M.A drafted the manuscript. All authors approved and contributed to the final manuscript.

## References

1. Breitbart M, Salamon P, Andresen B, Mahaffy JM, Segall AM, et al. Genomic analysis of uncultured marine viral communities. Proceedings of the National Academy of Sciences of the United States of America: Proc Natl Acad Sci U S A; 2002. p. 14250–14255.

2. Hurwitz BL, Hallam SJ, Sullivan MB. Metabolic reprogramming by viruses in the sunlit and dark ocean. Genome Biology: BioMed Central; 2013. p. R123.

3. Anantharaman K, Duhaime MB, Breier JA, Wendt KA, Toner BM et al. Sulfur oxidation genes in diverse deep-sea viruses. Science: American Association for the Advancement of Science; 2014. p. 757–760.

4. Roux S, Brum JR, Dutilh BE, Sunagawa S, Duhaime MB et al. Ecogenomics and potential biogeochemical impacts of globally abundant ocean viruses. Nature: Nature Publishing Group; 2016. p. 689-693.

5. Clooney AG, Sutton TDS, Shkoporov AN, Holohan RK, Daly KM et al. Whole-Virome Analysis Sheds Light on Viral Dark Matter in Inflammatory Bowel Disease. Cell Host & Microbe 2019. p. 764–778.e765.

6. Shkoporov AN, Clooney AG, Sutton TDS, Ryan FJ, Daly KM et al. The human gut virome is highly diverse, stable, and individual specific. Cell Host and Microbe: Elsevier Inc.; 2019. p. 527–541.e525.

7. Michniewski S, Rihtman B, Cook R, Jones MA, Wilson WH et al. A new family of “megaphages” abundant in the marine environment. ISME Communications 2021 1:1: Nature Publishing Group; 2021. p. 1-4.

8. Devoto AE, Santini JM, Olm MR, Anantharaman K, Munk P et al. Megaphages infect Prevotella and variants are widespread in gut microbiomes. Nature Microbiology 2019.

9. Adriaenssens EM, Roux S, Brister JR, Karsch-Mizrachi I, Kuhn JH et al. Guidelines for public database submission of uncultivated virus genome sequences for taxonomic classification. Nature Biotechnology 2023.

10. Simmonds P, Adams MJ, Benk M, Breitbart M, Brister JR, et al. Virus taxonomy in the age of metagenomics. Nature Reviews Microbiology 2017;15(3):161–168.

11. Simmonds P, Adriaenssens EM, Zerbini FM, Abrescia NGA, Aiewsakun P et al. Four principles to establish a universal virus taxonomy. PLOS Biology 2023;21(2):e3001922.

12. Bertrand D, Shaw J, Kalathiyappan M, Ng AHQ, Kumar MS et al. Hybrid metagenomic assembly enables high-resolution analysis of resistance determinants and mobile elements in human microbiomes. Nature Biotechnology 2019;37(8):937–944.

13. Tourancheau A, Mead EA, Zhang X-S, Fang G. Discovering multiple types of DNA methylation from bacteria and microbiome using nanopore sequencing. Nature Methods 2021;18(5):491–498.

14. Sereika M, Kirkegaard RH, Karst SM, Michaelsen TY, Sørensen EA et al. Oxford Nanopore R10.4 long-read sequencing enables the generation of near-finished bacterial genomes from pure cultures and metagenomes without short-read or reference polishing. Nature Methods 2022;19(7):823–826.

15. Kolmogorov M, Bickhart DM, Behsaz B, Gurevich A, Rayko M et al. metaFlye: scalable long-read metagenome assembly using repeat graphs. Nature methods: Nat Methods; 2020. p. 1103–1110.

16. Bickhart DM, Kolmogorov M, Tseng E, Portik DM, Korobeynikov A et al. Generating lineage-resolved, complete metagenome-assembled genomes from complex microbial communities. Nature Biotechnology 2022;40(5):711–719.

17. Feng X, Cheng H, Portik D, Li H. Metagenome assembly of high-fidelity long reads with hifiasm-meta. Nature Methods 2022;19(6):671–674.

18. Zhao Y, Temperton B, Thrash JC, Schwalbach MS, Vergin KL et al. Abundant SAR11 viruses in the ocean. Nature: Nature Publishing Group; 2013. p. 357-360.

19. Martinez-Hernandez F, Fornas Ò, Lluesma Gomez M, Garcia-Heredia I, Maestre- Carballa L et al. Single-cell genomics uncover Pelagibacter as the putative host of the extremely abundant uncultured 37-F6 viral population in the ocean. The ISME journal. 2018/09/18 ed: Nature Publishing Group UK; 2019. p. 232-236.

20. Olson ND, Treangen TJ, Hill CM, Cepeda-Espinoza V, Ghurye J et al. Metagenomic assembly through the lens of validation: recent advances in assessing and improving the quality of genomes assembled from metagenomes. Briefings in bioinformatics: Oxford University Press; 2019. p. 1140-1150.

21. Temperton B, Giovannoni SJ. Metagenomics: microbial diversity through a scratched lens. Curr Opin Microbiol 2012;15(5):605–612.

22. Mizuno CM, Ghai R, Rodriguez-Valera F. Evidence for metaviromic islands in marine phages. Frontiers in Microbiology: Frontiers Research Foundation; 2014.

23. Roux S, Emerson JB, Eloe-Fadrosh EA, Sullivan MB. Benchmarking viromics: An in silico evaluation of metagenome-enabled estimates of viral community composition and diversity. PeerJ 2017. p. e3817.

24. Cook R, Hooton S, Trivedi U, King L, Dodd CER et al. Hybrid assembly of an agricultural slurry virome reveals a diverse and stable community with the potential to alter the metabolism and virulence of veterinary pathogens. Microbiome 2021. p. 65.

25. Zablocki O, Michelsen M, Burris M, Solonenko N, Warwick-Dugdale J et al. VirION2: A shortand long-read sequencing and informatics workflow to study the genomic diversity of viruses in nature. PeerJ: PeerJ Inc.; 2021. p. e11088.

26. Warwick-Dugdale J, Solonenko N, Moore K, Chittick L, Gregory AC et al. Long-read viral metagenomics captures abundant and microdiverse viral populations and their niche- defining genomic islands. PeerJ 2019.

27. Elek CKA, Brown TL, Le Viet T, Evans R, Baker DJ et al. A hybrid and poly-polish workflow for the complete and accurate assembly of phage genomes: a case study of ten przondoviruses. Microbial Genomics 2023;9(7).

28. Wick RR, Judd LM, Holt KE. Assembling the perfect bacterial genome using Oxford Nanopore and Illumina sequencing. PLoS Comput Biol 2023;19(3):e1010905.

29. Buck D, Weirather JL, de Cesare M, Wang Y, Piazza P, et al. Comprehensive comparison of Pacific Biosciences and Oxford Nanopore Technologies and their applications to transcriptome analysis. F1000Research: Faculty of 1000 Ltd; 2017.

30. Watson M, Warr A. Errors in long-read assemblies can critically affect protein prediction. Nature Biotechnology 2019. p. 124–126.

31. Delahaye C, Nicolas J. Sequencing DNA with nanopores: Troubles and biases. PLOS ONE 2021;16(10):e0257521.

32. Kieft K, Zhou Z, Anantharaman K. VIBRANT: automated recovery, annotation and curation of microbial viruses, and evaluation of viral community function from genomic sequences. Microbiome: Cold Spring Harbor Laboratory; 2020. p. 90.

33. Cook R, Brown N, Rihtman B, Michniewski S, Redgwell T et al. The long and short of it: Benchmarking viromics using Illumina, Nanopore and PacBio sequencing technologies. bioRxiv 2023:2023.2002.2012.527533.

34. Beaulaurier J, Luo E, Eppley JM, Uyl PD, Dai X et al. Assembly-free single-molecule sequencing recovers complete virus genomes from natural microbial communities. Genome Research: Cold Spring Harbor Laboratory Press; 2020. p. 437-446.

35. Yilmaz S, Allgaier M, Hugenholtz P. Multiple displacement amplification compromises quantitative analysis of metagenomes. Nature methods: Nat Methods; 2010. p. 943–944.

36. Marine R, McCarren C, Vorrasane V, Nasko D, Crowgey E et al. Caught in the middle with multiple displacement amplification: The myth of pooling for avoiding multiple displacement amplification bias in a metagenome. Microbiome: BioMed Central Ltd.; 2014. p. 1–8.

37. Kim KH, Bae JW. Amplification Methods Bias Metagenomic Libraries of Uncultured Single-Stranded and Double-Stranded DNA Viruses. Applied and Environmental Microbiology: American Society for Microbiology (ASM); 2011. p. 7663.

38. Hsieh S-Y, Tariq MA, Telatin A, Ansorge R, Adriaenssens EM et al. Comparison of PCR versus PCR-Free DNA Library Preparation for Characterising the Human Faecal Virome. Viruses 2021;13(10):2093.

39. Roux S, Trubl G, Goudeau D, Nath N, Couradeau E et al. Optimizing de novo genome assembly from PCR-amplified metagenomes. PeerJ 2019;7:e6902.

40. De Coster W, Rademakers R. NanoPack2: population-scale evaluation of long-read sequencing data. Bioinformatics 2023;39(5).

41. Telatin A, Fariselli P, Birolo G. SeqFu: A Suite of Utilities for the Robust and Reproducible Manipulation of Sequence Files. Bioengineering 2021;8(5):59.

42. Li D, Luo R, Liu CM, Leung CM, Ting HF et al. MEGAHIT v1.0: A fast and scalable metagenome assembler driven by advanced methodologies and community practices. Methods: Academic Press Inc.; 2016. p. 3–11.

43. Langmead B, Salzberg SL. Fast gapped-read alignment with Bowtie 2. Nature methods: NIH Public Access; 2012. p. 357.

44. Kieft K, Adams A, Salamzade R, Kalan L, Anantharaman K. vRhyme enables binning of viral genomes from metagenomes. Nucleic Acids Research: Oxford University Press (OUP); 2022.

45. Nissen JN, Johansen J, Allesøe RL, Sønderby CK, Armenteros JJA et al. Improved metagenome binning and assembly using deep variational autoencoders. Nature Biotechnology 2021;39(5):555–560.

46. Vijini M, Michael JR, Bhavya P, Sarah KG, Susanna RG, et al. Phables: from fragmented assemblies to high-quality bacteriophage genomes. bioRxiv 2023:2023.2004.2004.535632.

47. Koren S, Walenz BP, Berlin K, Miller JR, Bergman NH et al. Canu: scalable and accurate long-read assembly via adaptive k-mer weighting and repeat separation. Genome Res 2017;27(5):722–736.

48. Ruan J, Li H. Fast and accurate long-read assembly with wtdbg2. Nature Methods: Nature Publishing Group; 2020. p. 155-158.

49. Vaser R, Šiki M. Time- and memory-efficient genome assembly with Raven. Nature Computational Science 2021;1(5):332–336.

50. Wick RR, Judd LM, Gorrie CL, Holt KE. Unicycler: Resolving bacterial genome assemblies from short and long sequencing reads. PLoS computational biology: PLoS Comput Biol; 2017.

51. Vaser R, Sovi I, Nagarajan N, Šiki M. Fast and accurate de novo genome assembly from long uncorrected reads. Genome Research: Genome Res; 2017. p. 737–746.

52. Li H. Minimap and miniasm: Fast mapping and de novo assembly for noisy long sequences. Bioinformatics: Oxford Academic; 2016. p. 2103–2110.

53. Li H. Minimap2: pairwise alignment for nucleotide sequences. Bioinformatics 2018. p. 3094–3100.

54. Li H. Aligning sequence reads, clone sequences and assembly contigs with BWA-MEM. arXiv preprint arXiv:13033997 2013.

55. Wick RR, Holt KE. Polypolish: Short-read polishing of long-read bacterial genome assemblies. PLOS Computational Biology: Public Library of Science; 2022. p. e1009802.

56. Camargo AP, Roux S, Schulz F, Babinski M, Xu Y et al. Identification of mobile genetic elements with geNomad. Nat Biotechnol 2023.

57. Nayfach S, Pedro Camargo A, Eloe-Fadrosh E, Roux S. CheckV: assessing the quality of metagenome-assembled viral genomes. bioRxiv: Cold Spring Harbor Laboratory; 2020. p. 2020.2005.2006.081778.

58. Hyatt D, Chen GL, LoCascio PF, Land ML, Larimer FW et al. Prodigal: Prokaryotic gene recognition and translation initiation site identification. BMC Bioinformatics: BioMed Central; 2010. p. 1–11.

59. Roux S, Adriaenssens EM, Dutilh BE, Koonin EV, Kropinski AM et al. Minimum Information about an Uncultivated Virus Genome (MIUViG). Nature biotechnology. 2018/12/17 ed: Nature Publishing Group US; 2019. p. 29-37.

60. Altschul SF, Gish W, Miller W, Myers EW, Lipman DJ. Basic local alignment search tool. Journal of Molecular Biology: Academic Press; 1990. p. 403–410.

61. Bin Jang H, Bolduc B, Zablocki O, Kuhn JH, Roux S et al. Taxonomic assignment of uncultivated prokaryotic virus genomes is enabled by gene-sharing networks. Nature Biotechnology: Nature Publishing Group; 2019. p. 632-639.

62. Cook R, Brown N, Redgwell T, Rihtman B, Barnes M et al. INfrastructure for a PHAge REference Database: Identification of Large-Scale Biases in the Current Collection of Cultured Phage Genomes. Phage: Cold Spring Harbor Laboratory; 2021. p. 214–223.

63. Gilchrist CLM, Chooi Y-H. clinker &amp; clustermap.js: automatic generation of gene cluster comparison figures. Bioinformatics 2021;37(16):2473–2475.

64. Team RC. R: A language and environment for statistical computing. Vienna: R Foundation for Statistical Computing; 2018.

65. Wickham H. Ggplot2: Elegant graphics for data analysis, 2 ed: Springer International Publishing; 2016.

66. Shannon P, Markiel A, Ozier O, Baliga NS, Wang JT et al. Cytoscape: A software environment for integrated models of biomolecular interaction networks. Genome Research: Cold Spring Harbor Laboratory Press; 2003. p. 2498-2504.

67. Conway JR, Lex A, Gehlenborg N. UpSetR: an R package for the visualization of intersecting sets and their properties. Bioinformatics 2017;33(18):2938–2940.

68. Sutton TDS, Clooney AG, Ryan FJ, Ross RP, Hill C. Choice of assembly software has a critical impact on virome characterisation. Microbiome: BioMed Central Ltd.; 2019. p. 1–15.

69. Gomez-Raya-Vilanova MV, Leskinen K, Bhattacharjee A, Virta P, Rosenqvist P et al. The DNA polymerase of bacteriophage YerA41 replicates its T-modified DNA in a primer- independent manner. Nucleic Acids Research 2022;50(7):3985–3997.

70. Rihtman B, Puxty RJ, Hapeshi A, Lee Y-J, Zhan Y et al. A new family of globally distributed lytic roseophages with unusual deoxythymidine to deoxyuridine substitution. Current Biology 2021;31(14):3199–3206.e3194.

71. Hutinet G, Kot W, Cui L, Hillebrand R, Balamkundu S et al. 7-Deazaguanine modifications protect phage DNA from host restriction systems. Nature Communications 2019;10(1):5442.

72. Bryson AL, Hwang Y, Sherrill-Mix S, Wu GD, Lewis JD et al. Covalent Modification of Bacteriophage T4 DNA Inhibits CRISPR-Cas9. mBio 2015;6(3):10.1128/mbio.00648-00615.

73. Flodman K, Tsai R, Xu MY, Corrêa IR, Jr., Copelas A et al. Type II Restriction of Bacteriophage DNA With 5hmdU-Derived Base Modifications. Front Microbiol 2019;10:584.

74. Thiaville JJ, Kellner SM, Yuan Y, Hutinet G, Thiaville PC et al. Novel genomic island modifies DNA with 7-deazaguanine derivatives. Proceedings of the National Academy of Sciences 2016;113(11):E1452–E1459.

75. Wick RR, Judd LM, Cerdeira LT, Hawkey J, Méric G et al. Trycycler: consensus long-read assemblies for bacterial genomes. Genome Biology 2021;22(1):266.

76. Kim CY, Ma J, Lee I. HiFi metagenomic sequencing enables assembly of accurate and complete genomes from human gut microbiota. Nature Communications 2022;13(1):6367.

77. Neri U, Wolf YI, Roux S, Camargo AP, Lee B et al. Expansion of the global RNA virome reveals diverse clades of bacteriophages. Cell 2022;185(21):4023–4037.e4018.

78. Conceição-Neto N, Yinda KC, Van Ranst M, Matthijnssens J. NetoVIR: Modular Approach to Customize Sample Preparation Procedures for Viral Metagenomics. Methods Mol Biol 2018;1838:85–95.

