## Supplementary figures and images for "Nanopore and Illumina Sequencing Reveal Different Viral Populations from Human Gut Samples"

### Supplementary Figure 1

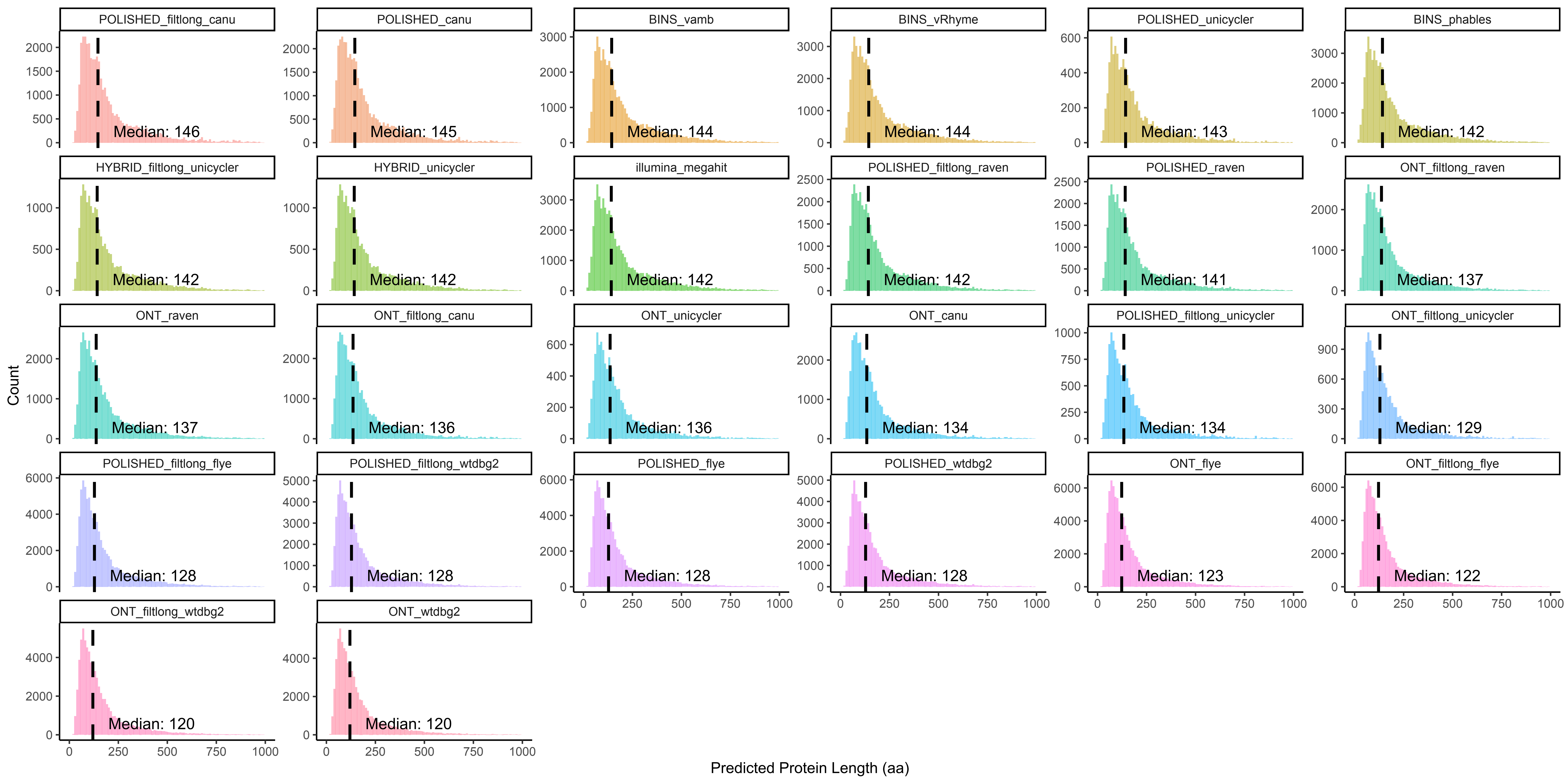
